# Rapid processing and quantitative evaluation of multicontrast EPImix scans for adaptive multimodal imaging

**DOI:** 10.1101/2021.02.12.430956

**Authors:** František Váša, Harriet Hobday, Ryan A. Stanyard, Richard E. Daws, Vincent Giampietro, Owen O’Daly, David J. Lythgoe, Jakob Seidlitz, Stefan Skare, Steven C. R. Williams, Andre F. Marquand, Robert Leech, James H. Cole

**Affiliations:** Department of Neuroimaging, Institute of Psychiatry, Psychology & Neuroscience, King’s College London, London, UK; Department of Forensic & Developmental Sciences, Institute of Psychiatry, Psychology & Neuroscience, King’s College London, London, UK; The Computational, Cognitive and Clinical Neuroimaging Laboratory, Department of Brain Sciences, Imperial College London, UK; Department of Child and Adolescent Psychiatry and Behavioral Science, Children’s Hospital of Philadelphia, Philadelphia, PA, USA; Department of Psychiatry, University of Pennsylvania, Philadelphia, PA, USA; Department of Neuroradiology, Karolinska University Hospital, Stockholm, Sweden; Department of Clinical Neuroscience, Karolinska Institutet, Stockholm, Sweden; Donders Institute for Brain, Cognition and Behavior, Radboud University Nijmegen, Nijmegen, The Netherlands; Department for Cognitive Neuroscience, Radboud University Medical Center Nijmegen, Nijmegen, The Netherlands; Department of Computer Science, Centre for Medical Image Computing, University College London, London, UK; Dementia Research Centre, Institute of Neurology, University College London, London, UK

**Author notes:** Corresponding author *Email address:* (František Váša). These authors have contributed equally (Robert Leech, James H. Cole).

**Keywords:** MRI, Identifiability, Fingerprinting, Structural Covariance, Morphometric Similarity Network, Reliability

## Abstract

Current neuroimaging acquisition and processing approaches tend to be optimised for quality rather than speed. However, rapid acquisition and processing of neuroimaging data can lead to novel neuroimaging paradigms, such as adaptive acquisition, where rapidly processed data is used to inform subsequent image acquisition steps. Here we first evaluate the impact of several processing steps on the processing time and quality of registration of manually labelled T_1_-weighted MRI scans. Subsequently, we apply the selected rapid processing pipeline both to rapidly acquired multicontrast EPImix scans of 95 participants (which include T_1_-FLAIR, T_2_, T_2_*, T_2_-FLAIR, DWI & ADC contrasts, acquired in ∼1 minute), as well as to slower, more standard single-contrast T_1_-weighted scans of a subset of 66 participants. We quantify the correspondence between EPImix and single-contrast T_1_-weighted scans, using correlations between voxels and regions of interest across participants, measures of within- and between-participant identifiability as well as regional structural covariance networks. Furthermore, we explore the use of EPImix for the rapid construction of morphometric similarity networks. Finally, we quantify the reliability of EPImix-derived data using test-retest scans of 10 participants. Our results demonstrate that quantitative information can be derived from a neuroimaging scan acquired and processed within minutes, which could further be used to implement adaptive multimodal imaging and tailor neuroimaging examinations to individual patients.

Graphical abstract.

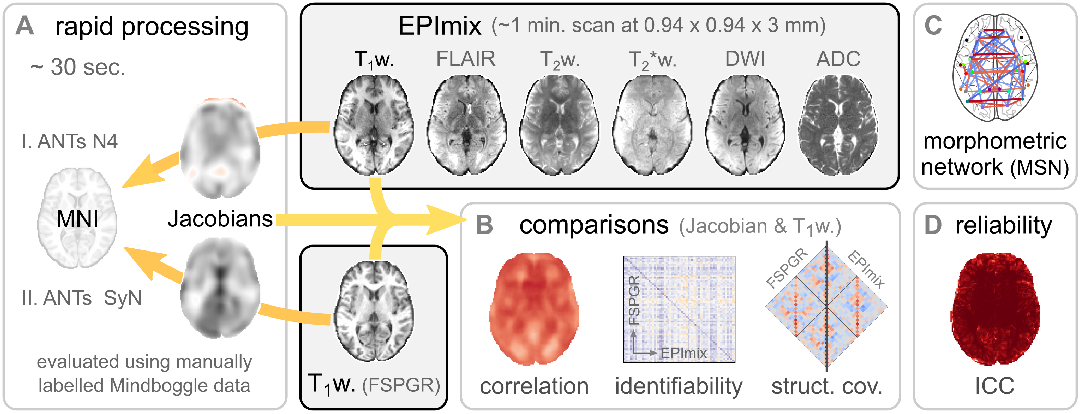

## Introduction

An MRI scanner can be used to acquire a range of different contrasts, which provide complementary information and are sensitive to different pathophysiologies (Cercignani and Bouyagoub, 2018). Currently, multimodal MRI scanning involves specifying a sequence of contrasts prior to data acquisition and in research contexts, acquiring the same sequence for each individual. In a clinical context, the selection of contrasts is guided by factors such as clinical history, cognitive and neurological examinations, and symptoms (Camprodon and Stern, 2013). However, the optimal sequence of contrasts and/or parameters for each contrast may depend on the anatomical or physiological abnormalities specific to the individual patient or be specific to a given pathology, and thus may not be known a priori.

As an alternative approach, it was recently proposed that data could be analysed as it is being acquired, with the near-real-time results used to determine subsequent acquisition steps (Cole et al., 2020). This approach was illustrated using three simulated scenarios, including (i) tailoring the resolution and/or field of view (FoV) of a structural scan to detect stroke, (ii) adaptively acquiring multimodal data to classify a known outcome variable using a decision tree, and (iii) adaptively searching across multiple MRI modalities using Bayesian optimisation to detect abnormality. However, adaptive acquisition is yet to be implemented practically. One prerequisite to progress beyond simulated scenarios (Cole et al., 2020) and implement adaptive acquisition in practice is the development of rapid analysis pipelines for multiple MRI modalities, enabling data to be processed in near-real-time.

We propose to capitalise on EPImix – a recently developed multicontrast sequence which acquires six contrasts (T_1_-FLAIR, T_2_, T_2_*, T_2_-FLAIR, DWI, ADC) in ~1 minute (Skare et al., 2018). EPImix is well suited to be the first sequence in an adaptive acquisition run, rapidly providing an overview of neuroanatomy across multiple contrasts. EPImix contrasts have previously been compared to high-quality, single-contrast sequences to evaluate their suitability for qualitative disease identification and categorisation by trained radiologists, and have shown comparable diagnostic performance to routine clinical brain MRI (Delgado et al., 2019; Ryu et al., 2020). However, there have been no quantitative comparisons of EPImix and corresponding single-contrast scans.

Here, we explore rapid image processing pipelines for the EPImix sequence, as well as for a single-contrast T_1_-weighted (T_1_-w) sequence, and use the rapidly processed scans to quantitatively compare EPImix and standard T_1_-w scans. We first optimise a rapid processing pipeline by evaluating the impact of several processing steps on the processing time and on the quality of registration of manually labelled scans, using openly available data with manual segmentations in both native and standard space (Klein and Tourville, 2012). Subsequently, we quantify, in several ways, the overlap between selected EPImix contrasts and corresponding single-contrast sequences. Finally, we demonstrate a novel quantitative application of the multicontrast EPImix sequence: the construction of morphometric similarity networks (MSNs; Seidlitz et al. (2018)).

## Methods

### Processing steps

While developing a rapid image processing pipeline, we considered the following factors to guide selection of steps:

- *Speed*: Faster processing was preferred. We measured speed in seconds. (Processing was run on an Apple Macbook Pro [2.2GHz Intel Core i7, 16Gb 1600MHz DDR3 RAM], with no other user processes running in parallel).
- *Quality*: Higher quality was preferred. We evaluated the quality of steps up to and including registration by quantifying overlap between source and target of manually labelled atlases (Klein et al., 2009) using the Mindboggle dataset (Klein and Tourville, 2012).
- *Automation*: Fewer quality control steps and resulting re-running of processing steps following manual interventions and/or changes of parameters were preferred.

For the processing steps considered for inclusion in the pipeline, see Table 1.

**Table 1:** Processing steps considered. For each step, we list the reason for consideration, the evaluated options and the algorithm used, including relevant references.

| Step | Reason | Options | Algorithm<br>(Reference(s)) |
| --- | --- | --- | --- |
| Downsampling | To save time<br>(& potentially help extraction) | 1 / 2 / 3 mm | ANTs *ResampleImageBySpacing*<br>(Avants et al., 2008, 2011) |
| Bias field correction | Commonly applied<br>to improve registration | yes / no | ANTs *N4BiasFieldCorrection*<br>(Tustison et al., 2010) |
| Brain extraction | To improve registration | yes / no | FSL *BET*<br>(Smith, 2002) |
| Registration | To evaluate deviation<br>from spatially normalised group | SyN / b-spline SyN | ANTs *antsRegistrationSyNQuick.sh*<br>(Avants et al., 2008, 2011) |
| Smoothing | To remove noise<br>in voxel-wise analyses | 2 / 4 / 6 mm FWHM | Python *nilearn nl.image.smooth_img*<br>(Abraham et al., 2014) |

For registration of scans to standard space, we used ANTs (Avants et al., 2008), due to its good performance in systematic evaluations of registration algorithms (Klein et al., 2009; Nazib et al., 2018; Bartel et al., 2019). We are aware that the combination of processing steps listed in Table 1 is by no means exhaustive, as different software suites could have been used for each step, potentially differing in speed and quality of processing; instead, the selected steps serve as a proof-of-principle evaluation of the proposed approach (see also *Discussion*).

We also note that, during application of these steps to single-contrast scans in real time, the data would not be downsampled; instead, it would directly be acquired at a reduced resolution. As the T_1_-w data used here was previously acquired at 1 mm isotropic resolution and is not being processed in real time, the downsampling step serves to simulate the effect of acquisition of the data at a lower resolution on processing. (We note that the EPImix acquisition is currently fixed to a 180 × 180 in-plane matrix size, so options for modifying the resolution during acquisition are limited. However, processing of EPImix scans with 0.9375 × 0.9375 × 3 mm resolution is sufficiently fast to be carried out without further downsampling; see below for details.)

### Evaluating the speed and quality of registrations

We evaluated the quality of registrations as well as the effect of any prior pre-processing steps using the Mindboggle dataset (Klein and Tourville, 2012), which contains T_1_-w scans of 101 healthy participants manually labelled according to the Desikan-Killiany-Tourville (DKT) protocol (31 cortical regions per hemisphere). The dataset contains both T_1_-w scans and manual DKT atlas labels in both native and MNI152 spaces. These manual labels have previously been used as a gold standard in evaluations of processing tools (e.g., Tustison et al., 2014; Velasco-Annis et al., 2017; Henschel et al., 2020). We used the non-skull-stripped T_1_-w scans as initial input into our processing pipelines as brain extraction is one of the processing steps under evaluation.

We first used the native space T_1_-w scan to estimate registration parameters to MNI152 space (following any optional pre-processing steps; Fig. 1A). Subsequently, we applied the registration step to the manual native space DKT atlas labels (Fig. 1B). Finally, we quantified the over-lap of the transformed atlas labels with the manual MNI152 space atlas labels using the Dice coefficient (Fig. 1C), equal to twice the number of overlapping voxels divided by the sum of the number of voxels in each set; for voxel sets {A} and {B}:

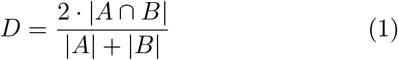

**Figure 1:**
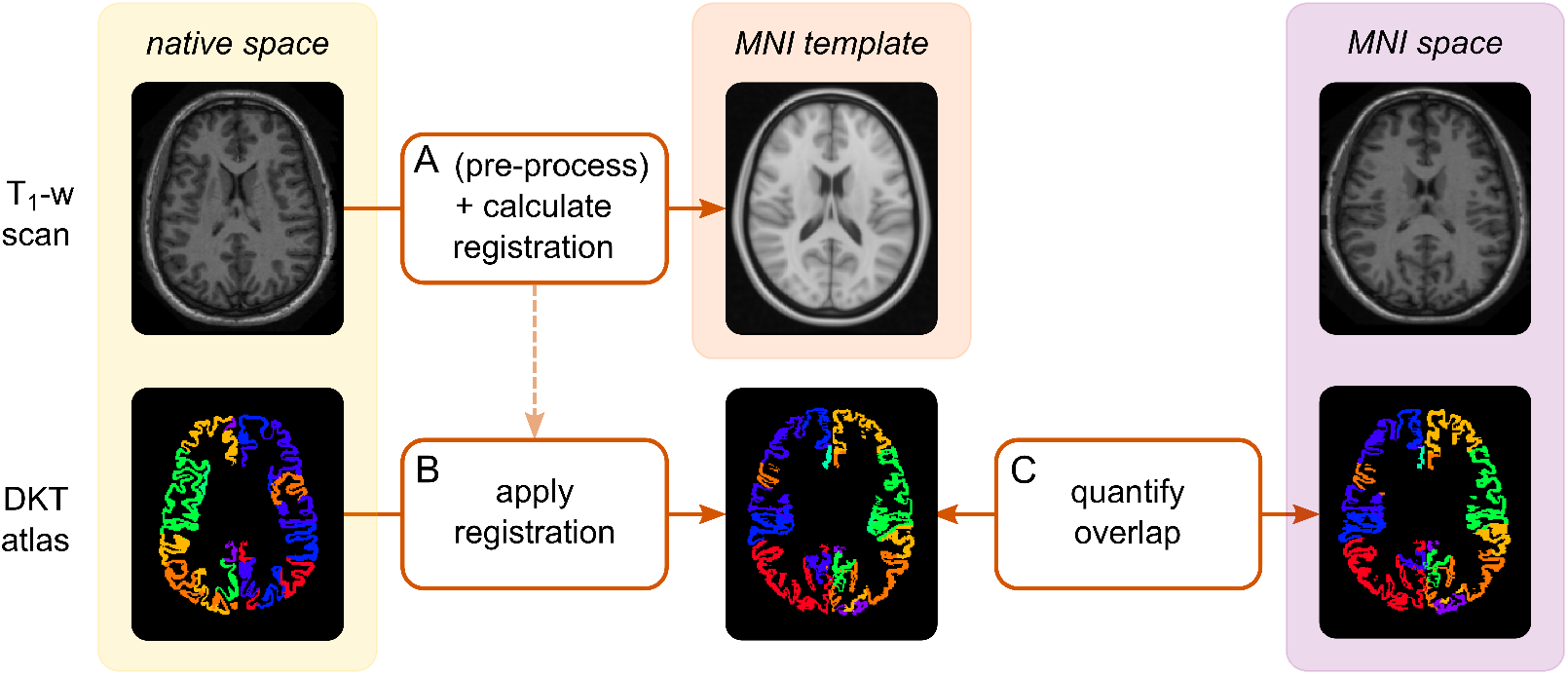
Using manual DKT atlas labels from the Mindboggle dataset to quantitatively evaluate the quality of registration (and pre-processing steps). A) The processing pipeline (up to and including registration) is applied to the native-space T_1_-w scan to transform it to MNI152 space and to estimate registration parameters. B) The registration (calculated in step A) is applied to the native-space DKT atlas. C) The Dice coefficient is used to quantify the overlap, in MNI152 space, between the atlas labels which have been transformed from native space (in step B) and the manual atlas labels released with the Mindboggle dataset.

We calculated the Dice coefficient both for all atlas regions across the brain at once, and for individual atlas regions.

We evaluated the above steps (Table 1) in a sequential manner, as follows. (As the specific selection of options evaluated at each step depended on results obtained in the previous step, we report the outcome of each step here; for details and reasons underlying our selection of each option, see *Results*. Unless otherwise specified, we used the ANTs SyN registration as implemented by default in *antsRegistrationSyNQuick.sh* as the main processing step.)

1. We first evaluated the effect of spatial resolution, including 1 mm (native Mindboggle data resolution), 2 mm and 3 mm isotropic. We downsampled both the T_1_-w scans and DKT atlases in both native and standard space, before applying the ANTs SyN algorithm for registration (Avants et al., 2008). (As previously noted, during application in real-time, single-contrast scans would be directly acquired at the desired resolution, rather than downsampled.)
2. Following selection of the resolution (2 mm), we considered the effect of bias field correction. We compared the quality and speed of ANTs SyN registration with and without ANTs N4 bias field correction (Tustison et al., 2010).
3. We next considered the impact of brain extraction on the output of the previous steps (2 mm with bias field correction). We compared the default non-skull-stripped registration to the application of FSL BET (Smith, 2002) for skull-stripping. (We used default BET parameters, except for the fractional intensity threshold which was set to 0.4 based on an initial test evaluation using a subset of scans.)
4. Finally, we applied ANTs spline-based SyN registration to the output of previous steps (2 mm with bias field correction & without skull-stripping) to compare speed and quality to standard ANTs SyN (Avants et al., 2008).

As a quality control step, the T_1_-w scan in MNI152 space (i.e., the output of Fig. 1A) was visually assessed to ensure a successful registration. For details of the settings used for each processing step in each evaluated pipeline, see Table 2.

**Table 2:**
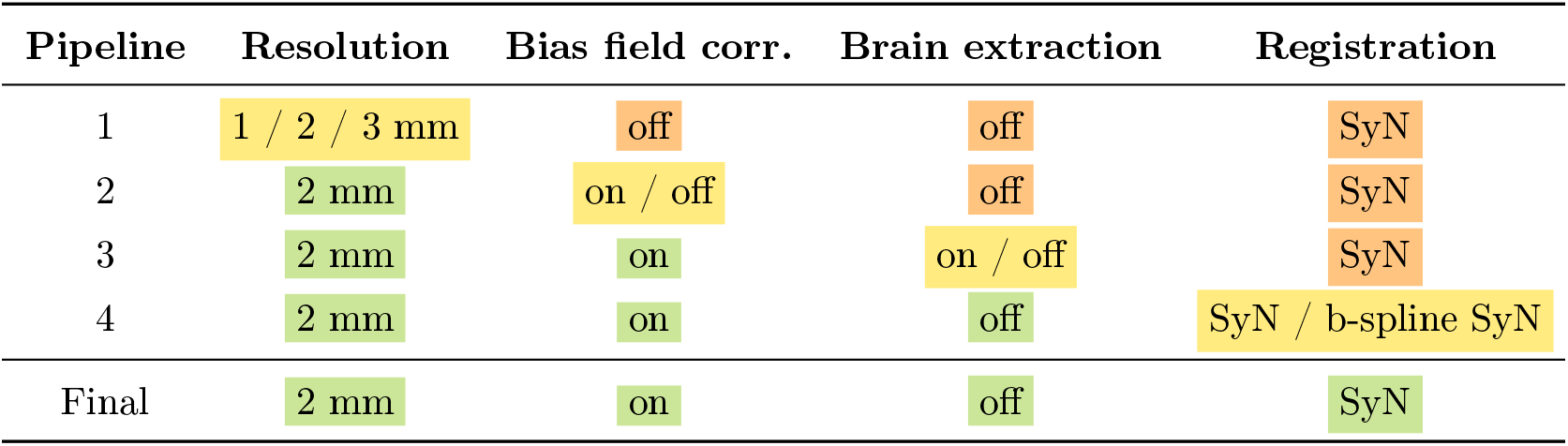
Evaluated pipelines. Settings used for each step while evaluating pipelines in a sequential manner. Cell colour indicates evaluation status: yellow cells indicate steps under evaluation, orange cells indicate steps not yet evaluated and green cells indicate evaluated steps, where an option has been selected.

| Pipeline | Resolution | Bias field corr. | Brain extraction | Registration |
| --- | --- | --- | --- | --- |
| 1 | 1 / 2 / 3 mm | off | off | SyN |
| 2 | 2 mm | on / off | off | SyN |
| 3 | 2 mm | on | on / off | SyN |
| 4 | 2 mm | on | off | SyN / b-spline SyN |
| Final | 2 mm | on | off | SyN |

We note that not all combinations of processing steps were systematically evaluated. Moreover, our aim was not to find the “optimal” processing pipeline, but rather to consider trade-offs in processing speed and quality, to identify a combination of processing steps which optimises both parameters (i.e., “good enough and fast enough”).

### Processing of EPImix and corresponding single-contrast T_1_-w scans

Following the selection of a rapid processing pipeline (2 mm scans with bias field correction and standard SyN registration, see also *Results*), we applied it to the EPImix scans, and corresponding T_1_-w single-contrast scans. We focused on the T_1_-w single-contrast sequence due to data availability.

We included scans collected on the same scanner (General Electric MR750 3.0T, Waukesha, WI) across three different studies conducted on healthy volunteers at the Centre for Neuroimaging Sciences, King’s College London’s Institute of Psychiatry, Psychology & Neuroscience. The studies received ethical approval from King’s College London’s Psychiatry, Nursing and Midwifery Research Ethics Committee (KCL Ethics References: HR-18/19-9268, HR-18/19-11058 and HR-19/20-14585). All participants gave written informed consent to take part in the study.

EPImix scans were collected from 95 participants (48 female, 47 male; age median [1st, 3rd Quartile] (Md [Q1,Q3]) = 25 [22,29] years; Supplementary Information (SI) Fig. S1), consisting of six contrasts (T2*, T2-FLAIR, T2, T_1_-FLAIR, DWI, ADC) acquired with the following parameters: T2*: TE = 28.5 ms, TR = 2430 ms; T2-FLAIR: TE = 113 ms, TR = 5797 ms, TI = 2751 ms; T2, DWI & ADC: TE = 90.5 ms, TR = 2272 ms; T_1_-FLAIR: TE = 16.5 ms, TR = 1300 ms, TI = 582 ms; flip angle = 90°, matrix size = 180 × 180, FoV = 240 mm, 32 slices, slice thickness = 3 mm, voxel resolution = 0.975 × 0.975 × 3 mm. The EPImix sequence includes an on-scanner motion correction step; the motion corrected images were used for further analyses. For further details regarding the EPImix sequence, see Skare et al. (2018). Additionally, for 10 participants, a second EPImix scan was acquired during the same session to investigate test-retest reliability.

Of the participants scanned with the EPImix sequence, 66 were additionally scanned, within the same session, with an IR-FSPGR T_1_-weighted sequence (33 female, 33 male; age Md [Q1,Q3] = 25 [23,29.75] years; SI Fig. S1). Of these, 12 were scans with the following parameters: TE = 3.172 ms, TR = 8.148 ms, TI = 450 ms, flip angle = 12°, matrix size = 256 × 256, FoV = 256 mm, 164 slices, slice thickness = 1 mm, voxel resolution 1 × 1 × 1 mm; and 54 were scans with the following parameters: TE = 3.016 ms, TR = 7.312 ms, TI = 400 ms, flip angle = 11°, matrix size = 256 × 256, FoV = 270 mm, 196 slices, slice thickness = 1.2 mm, voxel resolution = 1.05 × 1.05 × 1.2 mm.

When applying the previously identified rapid processing pipeline to EPImix scans, we omitted the downsampling step (to 2 mm isotropic resolution), as options for modifying the EPImix voxel resolution of 0.975 × 0.975 × 3 mm during acquisition are limited, and the “native” EPImix resolution resulted in sufficiently rapid processing (Md [Q1,Q3] = 32 [31,33] s across participants; see also Fig. 3). Instead, we registered EPImix T_1_-w scans directly to a 2 mm isotropic MNI template, and subsequently applied the same transformation to the remaining EPImix contrasts. Furthermore, following registration of the single-contrast and EPImix T_1_-w scans to MNI space, we extracted the logarithm of the Jacobian determinant of the ANTs SyN transform (combining the affine and non-linear warp components) to serve as an additional quantitative comparison of EPImix and corresponding single-contrast acquisitions (henceforth referred to as log-Jacobian).

**Figure 3:**
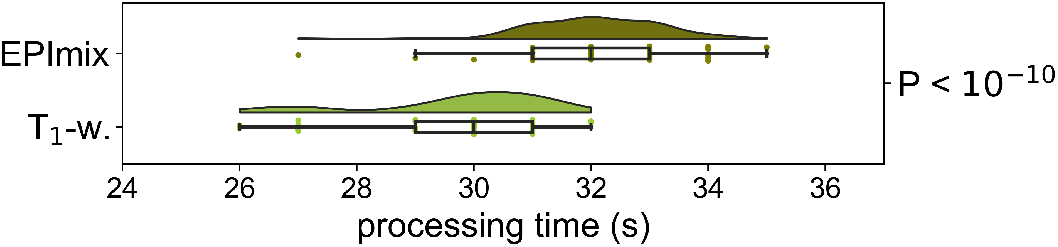
Processing time for EPImix and single-contrast T_1_-w scans. The P-value corresponds to the Mann-Whitney U test.

### Effects of resolution and spatial smoothness on the correspondence between EPImix and single-contrast T_1_-w scans

To evaluate the impact of spatial resolution, we down-sampled voxelwise data within regions of interest (ROIs). We used the 360 ROI atlas constructed by Glasser et al. (2016), as well as a downsampled version thereof, constructed by aggregating contiguous regions into 44 larger groups, as described in the Supplementary Information for Glasser et al. (2016). For details, see SI Fig. S2.

Due to the reduced FoV of EPImix scans, resulting in missing portions of the inferior temporal and/or superior parietal lobe in certain participants, we only included voxels present (i.e., non-zero) in at least 80% of EPImix scans in voxelwise analyses (i.e., 76/95 participants). For regional analyses, we only included regions of interest where at least 80% of voxels contained non-zero values in at least 80% of participants. This resulted in analyses using 297/360 regions from the Glasser et al., 2016 atlas, and 32/44 regions from its downsampled version. For details, see SI Fig. S3.

Regional values were generated by calculating the median values of unsmoothed voxel-wise EPImix contrasts, single-contrast T_1_-w scans and log-Jacobians within atlas masks registered to the same MNI space, excluding zero-valued voxels. We subsequently performed analyses at the spatial resolution of voxels (both spatially smoothed and unsmoothed), 297 and 32 ROIs, as described below. Additionally, voxel-wise analyses were performed and/or visualised using voxels within a mask defined by the MNI brain (dilated once), as well as cortical grey matter (GM) voxels (defined as voxels belonging to one of the regions of the cortical MMP atlases used).

Furthermore, to evaluate the impact of spatial smoothness on the correspondence between EPImix and corresponding single-contrast scans, we smoothed voxelwise EPImix and single-contrast T_1_-w scans using three different Gaussian kernels – 2, 4 and 6 mm full-width at half-maximum (FWHM; using Python *nilearn*; Abraham et al. 2014).

### Correspondence between EPImix and single-contrast scans

We quantified correspondence between matching EPImix and single-contrast T_1_-w scans in several ways. (All instances of correlation refer to Spearman’s correlation coefficient *ρ*.)

To evaluate the extent of spatial correspondence be-tween EPImix and single-contrast scans, we correlated corresponding log-Jacobians and T_1_-w intensities at the voxel and ROI level, across subjects.

Further, to determine whether the correspondence be-tween matching EPImix and single-contrast modalities is higher within than between participants, we calculated measures of “differential identifiability” (Amico and Goñi, 2018). This is defined as the median correlation of participants’ scans from one modality to their own scan from the other modality (i.e., within-participant correlation; *ρ*_*within*_), minus the median correlation between modalities of non-corresponding participants (i.e., between-participant correlation; *ρ*_*between*_):

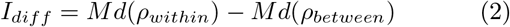

We additionally defined an individual index of differential identifiability, as the fraction of times that between-subject scan correlations are smaller than within-subject scan correlations. We calculated this measure twice for each participant and spatial resolution, to quantify both the individual identifiability of a single-contrast T_1_-w scan relative to EPImix T_1_-w scans, and of an EPImix T_1_-w scan relative to single-contrast T_1_-w scans. This individual measure of identifiability is related to discriminability, as defined by Bridgeford et al. (2020).

When correlating values at the regional level, we used a spatial permutation test to construct realistic null models of spatial correspondence. Permutations were generated by randomly rotating a projection of ROI centroids on the (FreeSurfer) sphere, before mapping rotated ROIs to the nearest unrotated ones. Mirrored rotations were applied to the contralateral hemisphere, resulting in a permutation which controls for spatial autocorrelation and hemispheric symmetry of regions (Váša et al., 2018; Alexander-Bloch et al., 2018; Markello and Misic, 2020). P-values for the correlation between two regional maps were obtained by comparing the empirical value of Spearman’s *ρ* to a null distribution of Spearman correlations, generated by correlating one of the empirical maps to a set of 10,000 spatially permuted versions of the other map; these “spin-test” P-values are referred to as P_*spin*_. Spin-test P-values were additionally corrected for multiple comparisons using the false discovery rate (FDR; Benjamini and Hochberg, 1995).

As a final comparison between contrasts, we used re-gional data from EPImix and single-contrast log-Jacobians as well as T_1_-w intensities to construct structural covariance matrices, by cross-correlating median regional values across subjects (Alexander-Bloch et al., 2013; Evans, 2013). We quantified correspondence between the upper triangular parts of the structural covariance matrices using correlation, and visualised networks from both modalities using (thresholded) network diagrams.

### EPImix morphometric similarity networks

We further explored the possibility of constructing morphometric similarity networks (MSNs; Seidlitz et al., 2018) from EPImix, by correlating regional contrast values between pairs of regions within subjects. We used seven maps per participant, including six EPImix contrasts as well as the log-Jacobian derived from transforming EPImix T_1_-w scans to MNI space. Regional values were normalised within each participant and contrast using the number of absolute deviations around the median, a non-parametric equivalent of the Z-score (Leys et al., 2013); for a vector of regional values *x*:

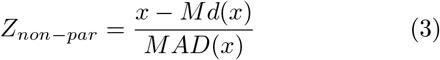

where *Md*() corresponds to the median, and *MAD*() to the median absolute deviation. Finally, normalised regional values were correlated using Spearman’s *ρ* across maps (contrasts), within participants, to create individual MSNs.

To explore the value of EPImix MSNs, we quantified the variance in participant age and sex explained by MSN edges using linear regression. The explained variance score was calculated within five-fold age-stratified cross-validation, with a resulting median value (across folds) calculated for each MSN edge.

### Test-retest reliability of EPImix scans

We quantified test-retest reliability of EPImix scans using 10 within-session test-retest EPImix scans. We quantified test-retest reliability using the intraclass correlation coefficient; specifically, we used the one-way random effects model for the consistency of single measurements, i.e., ICC(3,1), hereafter referred to as ICC (Chen et al., 2018). We calculated the ICC using voxel-wise data, ROI-averaged data and MSN edges.

## Results

### Evaluation of a rapid processing pipeline

We sequentially evaluated the impact of four processing steps on the spped and quality of registration, using the Mindboggle-101 dataset (Klein and Tourville, 2012). At each step, we recorded the processing time and the quality of overlap (between our custom registrations of DKT atlas labels and manual labels released with the Mindboggle dataset) using the Dice coefficient.

We first evaluated the impact of spatial resolution of the data. An isotropic resolution of 1 mm results in the most accurate registration, but is potentially too slow to be run in real-time (processing time Md [Q_1_,Q_3_] = 129 [127,131] s). The processing of the images with 2 mm isotropic resolution is sufficiently fast (Md [Q_1_,Q_3_] = 18 [18,19] s) and was therefore chosen (Fig. 2A). We next inspected the impact of bias field correction (on 2 mm isotropic resolution scans), using the ANTs N4 algorithm. We found that bias field correction improved registration quality at a relatively low time cost (Md [Q_1_,Q_3_] = 24 [24,25] s) and was therefore included as a processing step (Fig. 2B). Subsequently, we explored the application of a brain extraction algorithm (to the 2 mm isotropic resolution scans following bias field correction) using FSL BET. Brain extraction results in a marginally faster registration (Md [Q_1_,Q_3_] = 21 [20,23] s), but with no gain in quality (Fig. 2C). Combined with the fact that brain extraction might fail and need to be re-run with alternative parameters, it was not included in our processing pipeline. Finally, we evaluated the use of ANTs b-spline SyN registration (instead of the “standard” ANTs SyN algorithm). This results in a noticeably slower registration, without a gain in quality (Md [Q_1_,Q_3_] = 41 [40,42] s); therefore, the standard ANTs algorithm was preferred (Fig. 2D).

**Figure 2:**
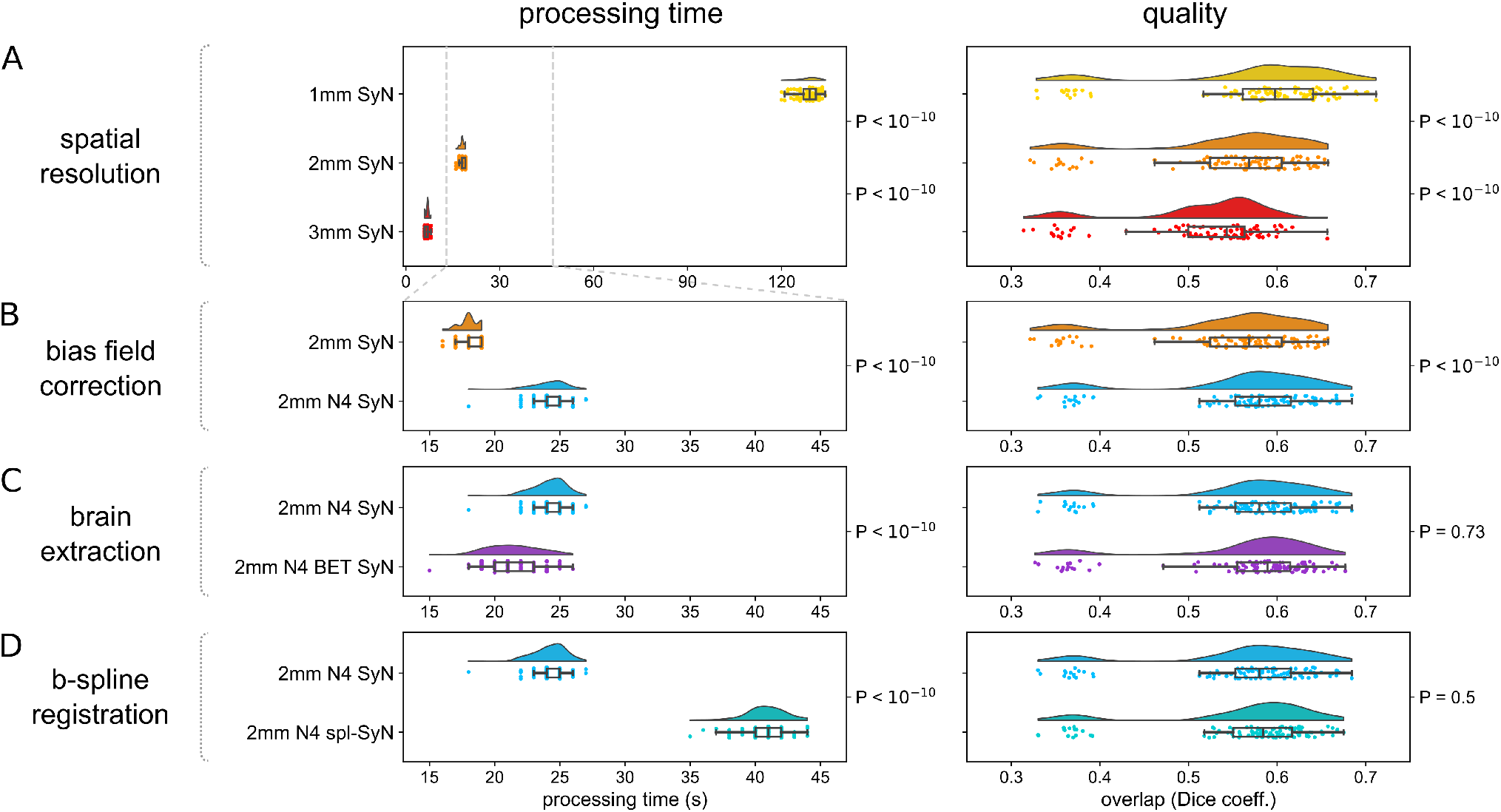
Evaluation of processing time and quality of registration using the Mindboggle dataset. The effect of four processing steps was evaluated sequentially; for each step, both processing time and quality were taken into account to select one of the options, before proceeding to the next step. P-values adjacent to neighbouring raincloud plots correspond to the (paired) Wilcoxon signed-rank test between corresponding data. A) Spatial resolution. B) Bias field correction. C) Brain extraction. D) B-spline SyN registration.

For each processing pipeline, we additionally calculated the Dice coefficient for individual regions of the DKT atlas. This showed a relatively spatially homogenous impact of processing steps on registration quality overall; for details, see SI Fig. S4.

We next applied the selected processing pipeline, consisting of ANTs N4 bias field correction and ANTs SyN registration, to EPImix and corresponding single-contrast T_1_-w scans (the EPImix scans were not downsampled but registered to a 2 mm isotropic MNI template brain directly; the single-contrast T_1_-w scans were downsampled to 2 mm isotropic resolution prior to registration). Application of the selected processing pipeline to EPImix and single-contrast T_1_-w scans resulted in rapid processing of both acquisitions (EPImix processing time Md [Q1,Q3] = 32 [31,33] s, single-contrast T_1_-w processing time Md [Q1,Q3] = 30 [29,31] s; Fig. 3).

### Correspondence between EPImix and single-contrast T_1_-weighted scans

We evaluated correspondence between EPImix and single-contrast scans using both log-Jacobians extracted from transformations of T_1_-w scans to MNI standard space, and T_1_-w scan intensities. In the main text, we report results of log-Jacobian comparisons as well as summary results for T_1_-w intensities; full details for comparisons of T_1_-w intensities are reported in the Supplementary Information.

We restricted analyses of EPImix and single-contrast T_1_-w scans to voxels with coverage in at least 80% participants (199’870/269’462 = 74.2% voxels in the MNI brain mask, and 64’370/78’247 = 82.3% voxels in the cortical GM mask), and regions where at least 80% voxels were non-zero in at least 80% participants (297/360 = 82.5% regions in the high-resolution MMP atlas, 32/44 = 72.7% regions in the low-resolution MMP atlas). For details regarding participant overlap at voxels and regions, see SI Fig. S3.

When evaluating correspondence between EPImix and single-contrast T_1_-w scans, we first calculated the correlation, across participants, of the log-Jacobian value at each voxel or region. All correlations were strong and positive, including *ρ* ≤ 0.96 at the voxel level (*ρ* ≤ 0.92 in the grey matter), and *ρ* ≤ 0.90 / 0.91 at the level of regions of the high / low resolution MMP atlas respectively (Fig. 4). Analogous comparisons using T_1_-w intensities yielded high voxel-wise correlations within the grey matter (Spearman’s *ρ* ≤ 0.80), with lower correlations in the white matter and within regions of interest of the high-resolution (*ρ* ≤ 0.41) and low resolution (*ρ* ≤ 0.30) MMP atlas (SI Fig. S5).

**Figure 4:**
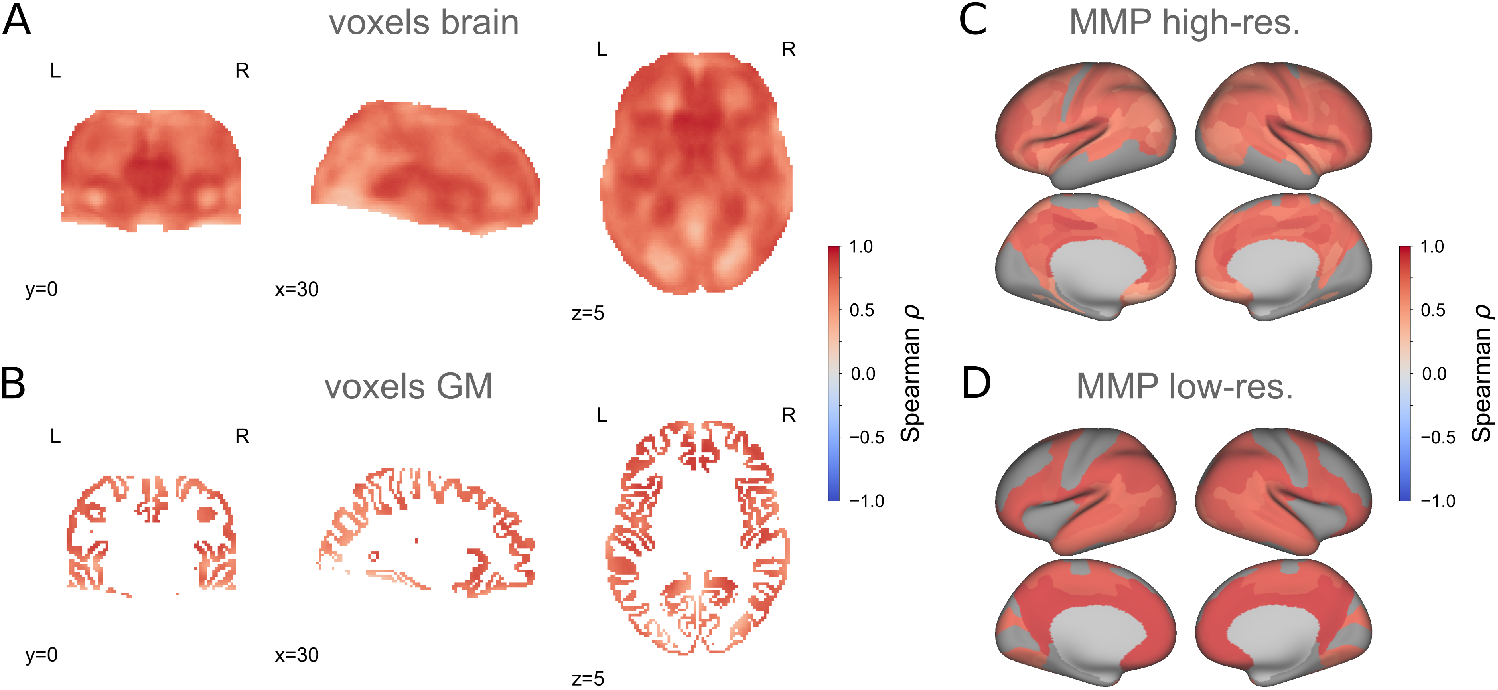
Local correspondence of log-Jacobians across participants. Spearman’s correlations between log-Jacobians of rapidly-processed T_1_-w scans from the EPImix sequence and a single-contrast acquisition, using data of 66 participants. Correlations are depicted: at the voxel level for A) the whole brain, and B) cortical grey matter, as well as within regions of interest of C) the high-resolution and D) the low-resolution MMP atlas. (At the regional level, median regional values were extracted prior to calculation of correlations for each region.)

We next quantified the within- and between-participant correspondence of EPImix and single-contrast data log-Jacobians (Fig. 5A). We calculated global identifiability, as the difference of the median between-participant correlation and median within-participant correlation (Fig. 5B; relevant parts of the correlation matrices are depicted in Fig. 5C). Differential identifiability was similar across types of data used, with highest identifiability at the level of low-resolution regions (I_*diff*_ = 0.49 – 0.16 = 0.33), closely followed by high-resolution regions (I_*diff*_ = 0.48 – 0.19 = 0.29), brain voxels (I_*diff*_ = 0.38 – 0.11 = 0.27) and finally GM voxels (I_*diff*_ = 0.40 – 0.14 = 0.26) (Fig. 5B). For regional data, we additionally used a null model relying on spherical “spin” permutation of cortical regions to account for spatial autocorrelation of the data when quantifying spatial correspondence between contrasts. Within the high-resolution atlas, 52/66 = 78.8% of within-participant correlations survived the FDR-corrected permutation test, compared to 406/4290 = 9.5% of between-participant correlations. Within the low-resolution atlas, no within- or between-participant correlations survived this thresholding procedure (Fig. 5A). Finally, we calculated individual-level identifiability, as the fraction of times that within-participant scan correlations are higher than between-participant scan correlations, using one of the contrasts as a reference (Fig. 5D). For example, identifiability of an individual EPImix T_1_-w scan is maximal (= 1) when the correlation between that scan and the same participants’ single-contrast T_1_-w scan is higher than all correlations to other participants’ single-contrast T_1_-w scans. Individual identifiability was highly similar when using EPImix T_1_-w scans and T_1_-w scans as reference. In contrast with global differential identifiability, individual participants were most identifiable at the level of brain voxels, with high individual identifiability at the level of GM voxels and high-resolution regions of interest as well; low-resolution regions had comparatively lower individual identifiability (Fig. 5D). Analogous analyses using T_1_-w scan intensities yielded highest differential and individual identifiability at the level of GM voxels (I_*diff*_ = 0.24; *Md*(ind. I_*diff*_) = 1), with lower correspondence at other spatial resolutions; for details, see SI Fig. S6.

**Figure 5:**
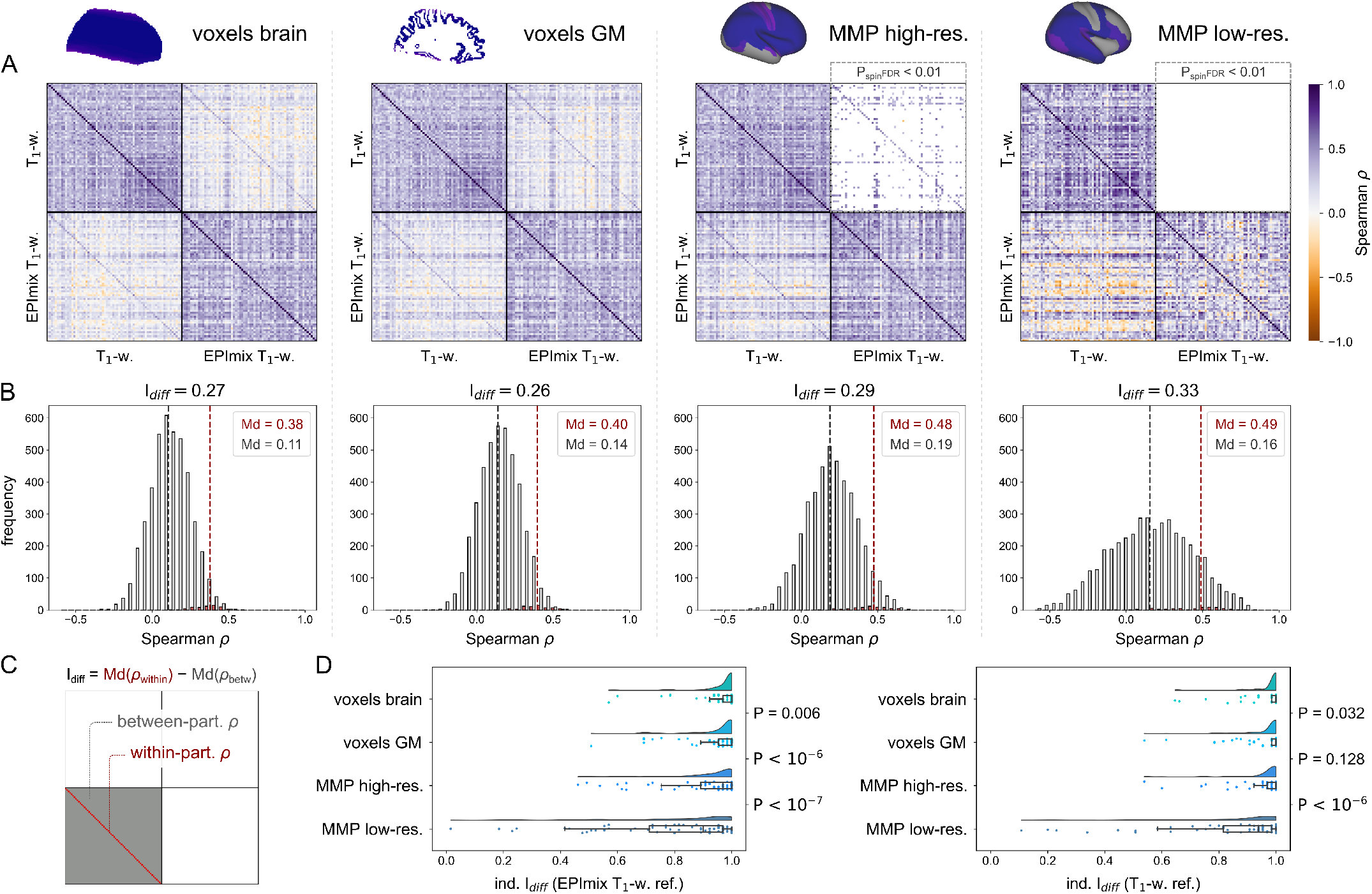
Participant identifiability across EPImix and single-contrast scans, using log-Jacobians. Between-participant correlations and identifiability were investigated using four types of data, at three spatial resolutions (columns in A-B, rows in D): all brain voxels, cortical grey matter voxels, regions of the high-resolution MMP atlas, and regions of the low-resolution MMP atlas. A) Spearman’s correlations between EPImix and single-contrast log-Jacobians, within and between participants. Cross-contrast correlations at the level of regions of interest were benchmarked using a null model controlling for contiguity and spatial autocorrelation (upper triangular blocks). B) Differential identifiability of contrasts, defined as the difference between the median within-participant correlation and the median between-participant correlation, as illustrated in C). D) Individual identifiability, defined as the fraction of times that the within-participant correlation is higher than between-participant correlations, either identifying the log-Jacobian of a single-contrast T_1_-w scan relative to log-Jacobians of EPImix T_1_-w scans (EPImix T_1_-w ref.), or vice-versa (T_1_-w ref.). P-values correspond to the Wilcoxon rank-sum test between neighbouring distributions.

We additionally inspected the effect of voxel-wise smoothing of T_1_-w intensity data, using 2, 4, and 6 mm FWHM kernels. The effect of smoothing was to reduce voxel-wise correspondence across subjects (i.e., the analogue of Fig. 4A,B), and to reduce differential identifiability due to a greater increase in the magnitude of between-participant correlations than within-participant correlations; for details, see SI Fig. S7.

For a summary of median within- and between-participant correspondence, and global and individual identifiability across spatial resolutions and types of data used (log-Jacobians and T_1_-w intensities), see Table 3.

**Table 3:** Identifiability of log-Jacobians and T_1_-w intensities at different spatial resolutions. From left to right, columns correspond to: median within-participant correlation, median between-participant correlation, global identifiability, individual identifiability with T_1_-w scans as reference, and individual identifiability with EPImix T_1_-w scans as reference. From top to bottom, blocks correspond to log-Jacobians and T_1_-w intensities (with rows corresponding to spatial resolution), as well as the effect of smoothing voxel-wise grey matter T_1_-w intensity maps (with rows corresponding to the width of the smoothing kernel).

| | | $Md(\rho_{with.})$ | $Md(\rho_{betw.})$ | $I_{diff}$ | ind. $I_{diff}$ $T_1$-w ref.<br>Md $[Q_1, Q_3]$ | ind. $I_{diff}$ Em ref.<br>Md $[Q_1, Q_3]$ |
| --- | --- | --- | --- | --- | --- | --- |
| **log-Jacobian** | voxels brain | 0.38 | 0.11 | 0.27 | 1.0 [0.98,1.0] | 1.0 [0.97,1.0] |
|  | voxels GM | 0.40 | 0.16 | 0.24 | 1.0 [0.98,1.0] | 1.0 [0.95,1.0] |
|  | MMP high-res. | 0.48 | 0.19 | 0.29 | 1.0 [0.97,1.0] | 0.97 [0.89,1.0] |
|  | MMP low-res. | 0.49 | 0.16 | 0.33 | 0.94 [0.82,0.98] | 0.90 [0.71,0.97] |
| **$T_1$-w intensity** | voxels brain | 0.62 | 0.50 | 0.12 | 1.0 [1.0,1.0] | 1.0 [1.0,1.0] |
|  | voxels GM | 0.47 | 0.23 | 0.24 | 1.0 [1.0,1.0] | 1.0 [1.0,1.0] |
|  | MMP high-res. | 0.61 | 0.43 | 0.19 | 1.0 [1.0,1.0] | 1.0 [1.0,1.0] |
|  | MMP low-res. | 0.78 | 0.66 | 0.12 | 0.91 [0.79,0.98] | 0.88 [0.72,0.97] |
| **$T_1$-w GM smoothed** | 2 mm FWHM | 0.50 | 0.26 | 0.24 | 1.0 [1.0,1.0] | 1.0 [1.0,1.0] |
|  | 4 mm FWHM | 0.59 | 0.38 | 0.21 | 1.0 [1.0,1.0] | 1.0 [1.0,1.0] |
|  | 6 mm FWHM | 0.64 | 0.49 | 0.16 | 1.0 [1.0,1.0] | 1.0 [1.0,1.0] |

### Correspondence of structural covariance networks across contrasts

We next inspected the correspondence between structural covariance networks, constructed by correlating log-Jacobians (or T_1_-w intensities) between all pairs of regions, across participants.

Structural covariance networks constructed using log-Jacobians exhibited similar hallmarks of organisation to structural covariance networks commonly constructed from regional cortical thickness or grey matter volume data, such as strong long-range inter-hemispheric correlations between homotopic regions (Fig. 6). The upper triangular parts of these matrices exhibited moderate correspondence between acquisitions, both for the high-resolution atlas (Spearman’s *ρ* = 0.41), and the low-resolution atlas (Spearman’s *ρ* = 0.46). Conversely, structural covariance networks constructed using T_1_-w scan intensities showed lower correspondence between acquisitions, particularly within the high-resolution atlas; this was likely due to unusually high short-range correlations clustered in the frontal cortex in the EPImix T_1_-w data (SI Fig. S8).

**Figure 6:**
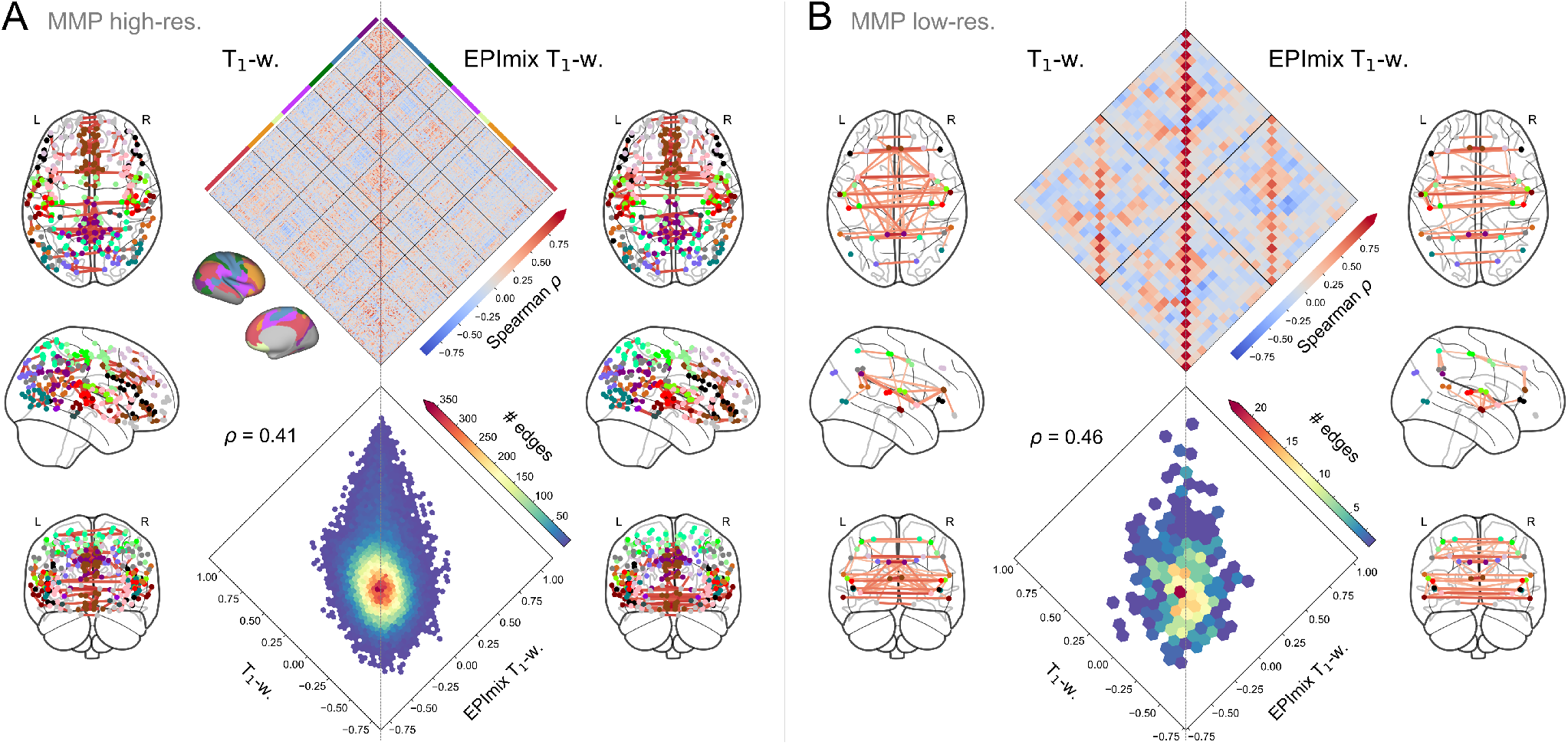
Structural covariance networks constructed from EPImix and single-contrast log-Jacobians. A) Structural covariance networks constructed using the high-resolution MMP atlas (297 regions). The diamond plot (top) is ordered according to regional membership of the 7 canonical intrinsic connectivity networks derived by Yeo et al. (2011). Network diagrams depict the strongest 0.3% correlations. B) Structural covariance networks constructed using the low-resolution MMP atlas (32 regions). Network diagrams depict the strongest 10% correlations.

### EPImix morphometric similarity networks

As a unique application of the multicontrast EPImix sequence, we explored the possibility of constructing morphometric similarity networks (MSNs; Seidlitz et al., 2018). We constructed individual MSNs by correlating nonparametrically normalised regional values of the six EPImix contrasts as well as the log-Jacobian (seven regional features in total), between all pairs of regions. For example, in the low-resolution MMP atlas, the left posterior opercular cortex showed high morphometric similarity to the left early auditory cortex (A1), but low similarity to the left posterior cingulate cortex (Fig. 7A). MSNs showed high positive correlations between homotopic pairs of regions (Fig. 7B).

**Figure 7:**
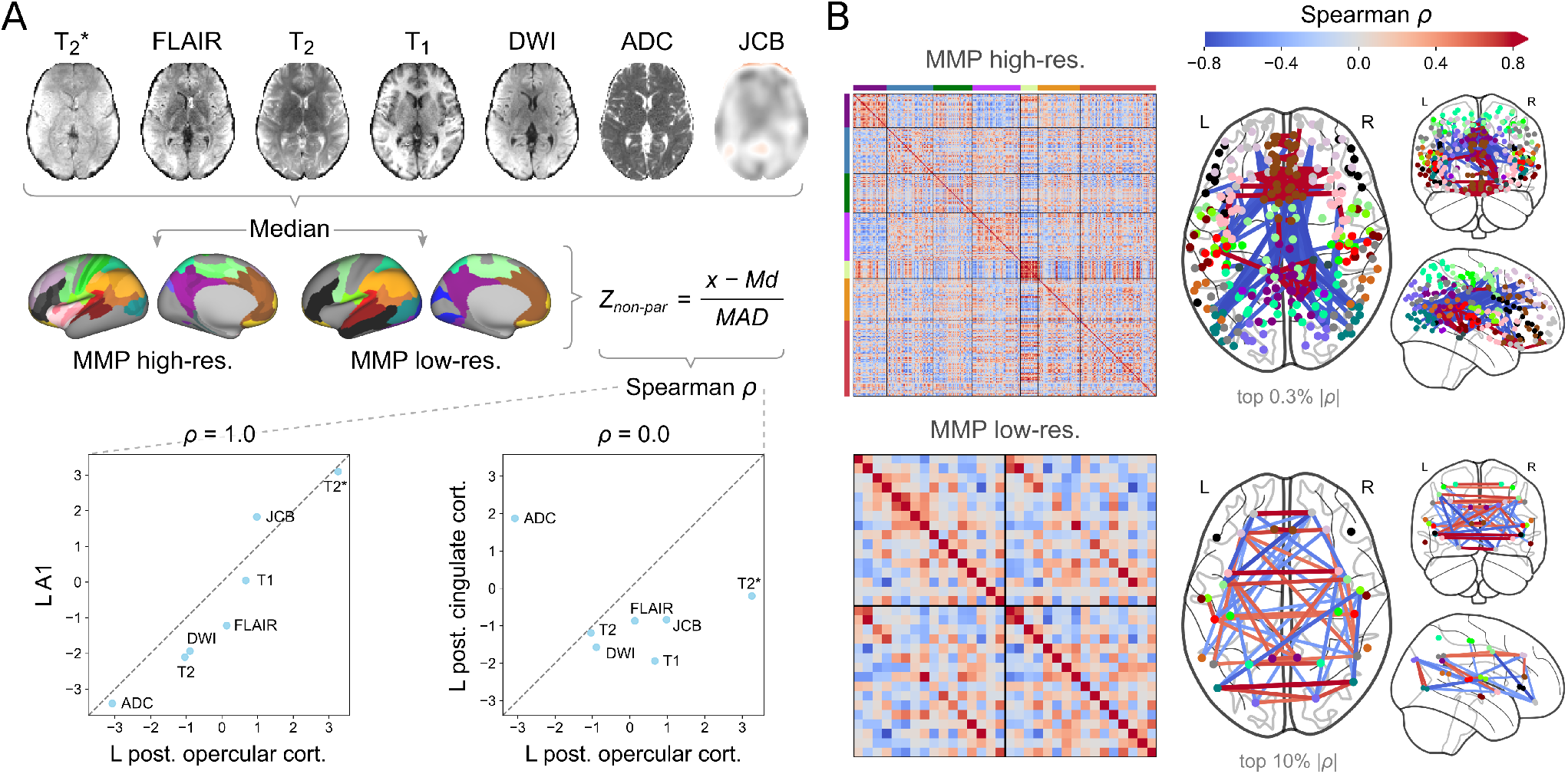
Morphometric similarity networks (MSNs) constructed from EPImix contrasts and log-Jacobians. A) MSN construction. Seven maps, including six EPImix contrasts and a log-Jacobian map (obtained from the warp of the T_1_-w contrast to MNI space) were used for network construction. Median values of each map within each region of a high-resolution and a low-resolution atlas were calculated, before normalisation (within participants, across regions) using a non-parametric equivalent of the Z-score (Md = Median, MAD = median absolute deviation). Two example correlations from the low-resolution atlas are shown: high morphometric similarity of left posterior opercular cortex to left early auditory cortex (A1), and low morphometric similarity to the left posterior cingulate cortex. B) Average MSNs (across participants), constructed using the high-resolution atlas (top; strongest 0.3% absolute correlations shown) and low-resolution atlas (bottom; strongest 10% absolute correlations shown).

We further explored the relationships of MSN edges to age and sex using all 95 participants with EPImix scans. At each edge of both the high-resolution and low-resolution MSN networks, we predicted edge strength (correlation) as a function of age and sex, using age-stratified five-fold cross-validation. We fitted each model to 80% (76/95) participants, and used the remaining 20% (19/95) participants to extract an explained variance score. Edge-wise median explained variance reached maximal values of 0.35 for high-resolution MSN edges, and 0.18 for low-resolution MSN edges.

### EPImix test-retest reliability

Finally, we quantified the test-retest reliability of rapidly processed EPImix contrasts, log-Jacobians and MSNs, using within-session test-retest data from 10 participants. Reliability was consistent across levels of spatial resolution, including voxels and regions of interest, and generally very high (Fig. 8 & Table 4). Among contrast maps, re-liability was lowest for the ADC; all other maps showed high reliability. MSN edges showed high reliability, with a few exceptions. For median and quartile ICC values, see Table 4.

**Figure 8:**
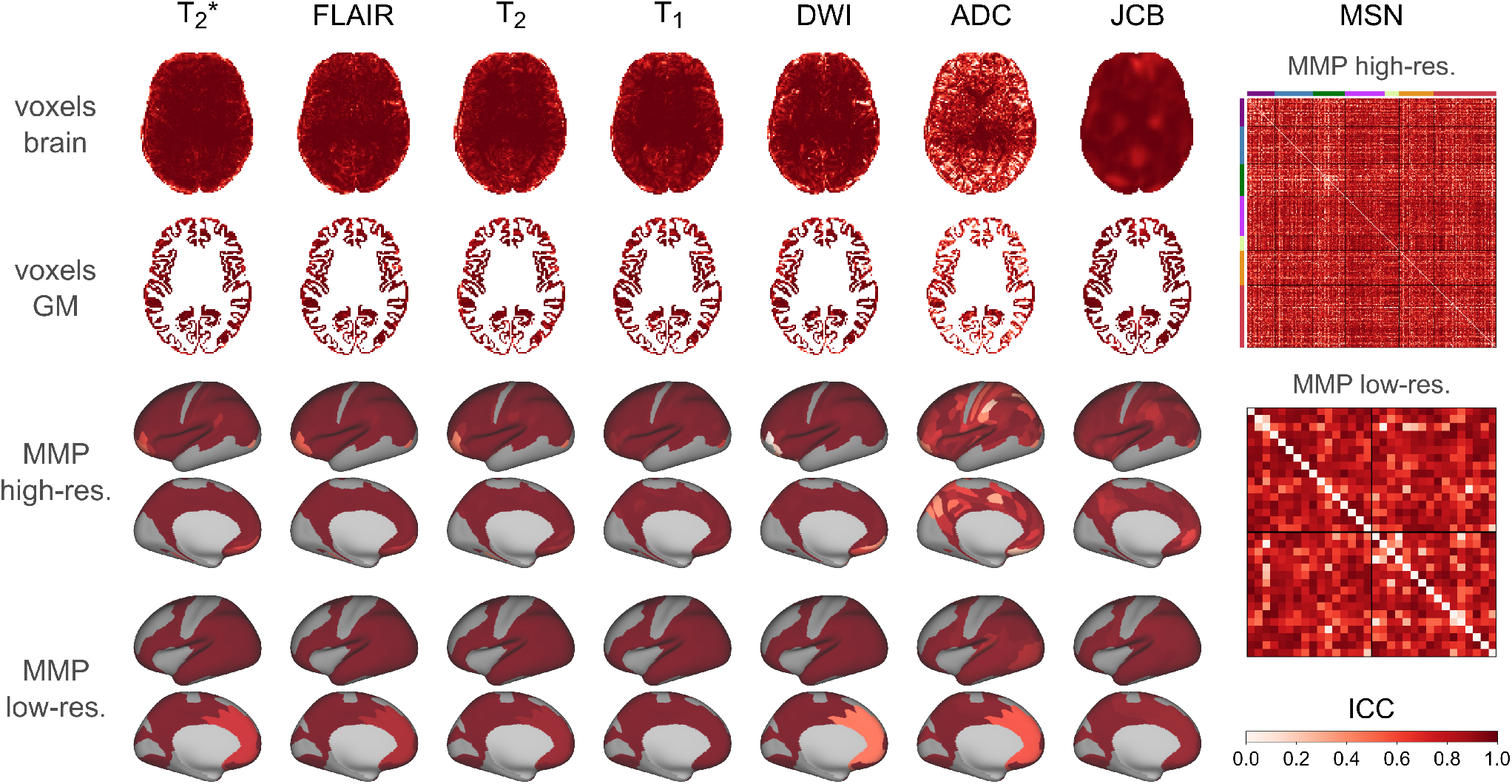
Test-retest reliability of rapidly-processed EPImix scans. Reliability was assessed using 10 within-session test-retest scans, for the six EPImix contrast and the log-Jacobian (JCB), at the level of voxels, and high- and low-resolution MMP atlases, as well as for morphometric similarity networks (MSNs) at both atlas resolutions. Reliability was quantified u sing the o ne-way random effects model for the consistency of single measurements, i.e., ICC(3,1).

**Table 4:** Test-retest reliability of values derived from 10 EPImix scans. Reliability was assessed for six EPImix contrasts as well as the log-Jacobian and MSNs, as Median [Q_1_,Q_3_] ICC(3,1).

| ICC: Md [Q<sub>1</sub>,Q<sub>3</sub>] | voxels brain | voxels GM | MMP high-res. | MMP low-res. |
| --- | --- | --- | --- | --- |
| T<sub>2</sub>* | 0.97 [0.93,0.99] | 0.97 [0.93,0.99] | 1.0 [0.99,1.0] | 1.0 [1.0,1.0] |
| (T<sub>2</sub>-)FLAIR | 0.97 [0.91,0.99] | 0.96 [0.89,0.98] | 1.0 [0.99,1.0] | 1.0 [1.0,1.0] |
| T<sub>2</sub> | 0.96 [0.88,0.99] | 0.94 [0.86,0.98] | 0.99 [0.98,1.0] | 1.0 [0.99,1.0] |
| T<sub>1</sub>(-FLAIR) | 0.96 [0.9,0.99] | 0.94 [0.88,0.97] | 0.99 [0.98,1.0] | 1.0 [0.99,1.0] |
| DWI | 0.96 [0.86,0.99] | 0.94 [0.80,0.98] | 0.99 [0.98,1.0] | 1.0 [0.99,1.0] |
| ADC | 0.78 [0.55,0.91] | 0.74 [0.48,0.89] | 0.91 [0.79,0.96] | 0.97 [0.93,0.99] |
| log-Jacobian | 0.96 [0.92,0.98] | 0.96 [0.92,0.98] | 0.97 [0.94,0.98] | 0.98 [0.97,0.99] |
| MSN | N/A | N/A | 0.81 [0.66,0.89] | 0.83 [0.69,0.90] |

## Discussion

### Rapid processing of MRI data

Using manually labelled scans from the Mindboggle dataset (Klein and Tourville, 2012), we first evaluated the impact of several processing steps on the processing time and quality of registration. The results informed our choice of “minimal” pre-processing pipeline, which included N4 bias field correction and ANTs SyN registration of non-skull-stripped scans at 2 mm isotropic resolution. This combination of processing steps resulted in very fast (< 1 minute on a typical computer) processing of both EPImix and single-contrast T_1_-w scans.

We further demonstrated that EPImix scans processed using our rapid pipeline showed high test-retest reliability. It would be interesting, in future, to repeat these analyses using test-retest data acquired during different sessions and/or at different scanner sites, for a more stringent test of the reliability and robustness of our rapid processing approaches (Chen et al., 2018).

We did not systematically evaluate all combinations of processing steps, in part due to the assumption that variation in registration quality (one of the underlying objective functions) will be relatively smooth along axes corresponding to each processing step (Lancaster et al., 2018); in other words, the interactions of the processing steps were assumed to be largely linear. Moreover, our evaluation was limited to a single algorithm per processing option. Use of other algorithms or software suites may have resulted in processing pipelines that run faster and/or result in higher quality registrations (Klein et al., 2009). In future, the multiverse of processing steps and algorithms could be more systematically and efficiently explored using adaptive methods, to simultaneously quantify the performance of different combinations of processing tools and identify optimal approaches (Dafflon et al., 2020).

Deep learning tools are another promising avenue for further optimisation of image processing. Such tools can substantially reduce the runtime of intensive processing steps (e.g., Henschel et al., 2020). Many of these tools require computationally expensive training on large datasets or are limited to specific applications, although new methods are being developed that are relevant to a broad range of tasks (Isensee et al., 2020). Another recent development is deep-learning methods trained on synthetic data, which promise to generate accurate segmentation (Billot et al., 2020) and registration (Hoffmann et al., 2020) in seconds, without the usual requirements of large empirical training datasets. The runtime/quality trade-off of these tools could be benchmarked using the tools presented here.

### Correspondence between EPImix and single-contrast scans

We quantified the correspondence between log-Jacobians and T_1_-w intensities derived from EPImix and single-contrast scans. Correspondence was generally high, across different spatial resolutions (including voxels and regions of interest) and higher within participants than between them, leading to high levels of participant identifiability. For log-Jacobians, global identifiability was highest for regions of the low-resolution MMP atlas, closely followed by other data resolutions; however, individual identifiability was similarly high at voxels and high-resolution regions, but lower for low-resolution regions. For T_1_-w intensities, both global and individual identifiability were highest using un-smoothed grey matter voxels. Taken together, these results suggest that grey-matter voxels and/or high-resolution regions of interest may be the optimal levels of spatial resolution for further analyses of similar data, for example in a real-time setting (Cole et al., 2020). An alternative approach would be to repeat analyses across spatial resolutions to generate a consensus result.

We note that between-contrast correspondence and identifiability were generally higher for log-Jacobians than for T_1_-w intensities. This could be due to the sensitivity of “raw” scan intensities to noise, including known variability due to factors such as scanner site or scan parameters (Shinohara et al., 2014). Methods have been proposed to harmonise intensities across scans, which involve matching distribution intensities within a reference section of tissue and accordingly adjusting intensities in other tissue classes (Shinohara et al., 2014; Fortin et al., 2016). However, such methods assume that variability within the reference tissue class is undesirable and thus preclude hypothesis testing in this tissue. The use of log-Jacobians provides an alternative avenue for quantitative analyses, which might be less affected by undesirable confounds.

As an additional means of comparing EPImix and single-contrast scans, we constructed structural covariance networks from regional log-Jacobians and T_1_-w intensities. One possible application of these networks is the calculation of individual contributions to group structural co-variance, as a single-contrast network biomarker of disease (Saggar et al., 2015). Given the modest between-contrast correspondence and apparently abnormal organisation of structural covariance networks derived from T_1_-w scan intensities, structural covariance networks constructed from regional log-Jacobian values are a more promising avenue for further work.

An alternative approach for deriving measures of brain connectivity from EPImix scans relies on morphometric similarity networks (MSNs; Seidlitz et al., 2018), constructed from within-participant correlations between regional morphometric features. Here we used six EPImix contrasts as well as the log-Jacobian to derive individual MSNs. MSNs have previously been constructed from 5-10 features, derived either from T_1_-w scans (often processed with FreeSurfer) or multiple contrasts (Seidlitz et al., 2018; King and Wood, 2020). Our rapidly-derived MSNs showcase the possibility of constructing individual multicontrast brain networks within minutes of participants entering the scanner.

We limited our comparison to EPImix and single-contrast T_1_-w scans (and corresponding log-Jacobians), due to lack of availability of high-resolution single-contrast data analogous to other EPImix contrasts for the same participants. Further work on developing rapid processing pipelines for additional sequences would be valuable, along with quantitative evaluations of within-participant correspondence between other EPImix contrasts and corresponding single-contrast scans. The methods used here to compare T_1_-w scans within participants can easily be translated to these other sequences, or further used to compare quantitative measures derived from other rapidly acquired scans. As an example, contrasts acquired on the recently developed low-field Hyperfine scanner (Sheth et al., 2020) could be compared to their high-resolution counterparts using the tools described herein.

### Towards adaptive imaging

The processing pipeline explored here is fast enough to be used while participants are still in the scanner, satisfying one of the key conditions for practical implementation of adaptive imaging (Cole et al., 2020). For the specific development of adaptive multimodal imaging, another requirement is a criterion for selecting which imaging modality or contrast to acquire next, based on hitherto acquired rapidly processed data.

One such criterion can be the selection of modalities predicted to show greatest deviations relative to large normative datasets (Marquand et al., 2019), based on previously acquired scans as well as knowledge of covariance across modalities. Thus, it would be valuable to use large multimodal normative datasets, such as CamCAN (Taylor et al., 2017), Human Connectome Project (Van Essen et al., 2012) or UK Biobank (Miller et al., 2016), to investigate both normative deviations of rapidly processed data across modalities as well as covariance between modalities across participants. In this context, repetition of analyses across spatial scales, such as voxels and regions of interest, could generate insights into the potential extent (and therefore nature) of abnormalities.

As a proof-of-concept of adaptive acquisition, we propose to use rapid processing and analysis of EPImix scans to determine which of the six contrasts show the greatest deviations from a normative population (Marquand et al., 2019). Subsequently, and while the participant is still in the scanner, these contrasts would be re-acquired at higher resolution using single-contrast sequences, in the order automatically determined by our rapid analysis algorithm. Subsequent “off-line” analyses could confirm the accuracy of the order of contrasts by extent of deviation from the norm, previously determined in real-time.

Practical implementation of adaptive multimodal imaging would enable personalised neuroimaging examinations of patients. This would potentially lead to reduced scanning time and cost and consequently greater patient comfort, as well as to a decreased likelihood of recalling patients for further examinations (Cole et al., 2020). An added benefit of adaptive methods in the research context is the reduced likelihood of questionable practices such as P-hacking or SHARKing (selecting hypotheses after results are known; Poldrack et al., 2017), due to the combination of data acquisition and analysis in a closed loop (Lorenz et al., 2017).

### Conclusion

In summary, we explored the impact of several rapid processing steps on the runtime and quality of registration, and used results to inform the choice of steps forming a minimal processing pipeline. Subsequently, we quantified the correspondence between rapidly processed multicontrast EPImix and single-contrast T_1_-w scans, demonstrating that substantial quantitative information can be reliably extracted from the EPImix sequence in minutes. Finally, we explored the use of EPImix for the rapid construction of morphometric similarity networks. Our work constitutes a step towards adaptive multimodal imaging, where real-time scan processing and analysis can inform tailoring of neuroimaging examinations to individual patients.

## Availability of code and data

All processing and analysis code is available on FV’s github, at https://github.com/frantisekvasa/epimix_rapid_processing. Mindboggle data is publicly available at https://osf.io/ydyxu/. Processed EPImix and single-contrast T_1_-w data, including contrast intensities and log-Jacobians at voxels of the MNI brain, is available at https://doi.org/10.6084/m9.figshare.13862729.

## Acknowledgments

RL & SCRW received support from the Wellcome/EPSRC Centre for Medical Engineering (WT 203148/Z/16/Z). FV, RL & SCRW would also like to acknowledge support from the Data to Early Diagnosis and Precision Medicine Industrial Strategy Challenge Fund, UK Research and Innovation (UKRI). We would also like to acknowledge support from NIHR Maudsley Biomedical Research Centre (BRC) NNA08. Finally, we thank Arno Klein and Jason Tourville for openly sharing the Mindboggle dataset.

