## Supplementary Information for "Rapid processing and quantitative evaluation of multicontrast EPImix scans for adaptive multimodal imaging"

### Correspondence between EPImix and single-contrast $T_1$-weighted scan intensities

Local correlations of $T_1$-w intensities were generally positive. At the voxel level, correlations were highest in the grey matter and cerebrospinal fluid (Spearman's $\rho \leq 0.80$), and lower in white matter (Fig. S5A,B). Within regions of interest of the MMP atlases, correlations were lower but predominantly positive, both at the high resolution ($\rho \leq 0.41$; Fig. S5C) and at the low resolution ($\rho \leq 0.30$; Fig. S5D).

We next quantified the within- and between-participant correspondence of EPImix and single-contrast data (Fig. S6A). We calculated global identifiability, as the difference of the median between-participant correlation and median within-participant correlation (Fig. S6B; relevant parts of the correlation matrices are depicted in Fig. S6C). Identifiability was low at the level of brain voxels ($I_{diff} = 0.62 - 0.50 = 0.12$), but considerably higher when correlating cortical GM voxels only ($I_{diff} = 0.47 - 0.23 = 0.24$). Averaging intensities within regions of interest led to increases in both within-participant and between-participant correlations, resulting in decreased identifiability – both for the high-resolution atlas ($I_{diff} = 0.61 - 0.43 = 0.19$) and the low-resolution atlas ($I_{diff} = 0.78 - 0.66 = 0.12$). For regional data, we additionally used a null model relying on spherical "spin" permutation of cortical regions to account for spatial autocorrelation of the data when quantifying spatial correspondence between contrasts. Within the high-resolution atlas, 66/66 = 100% of within-participant correlations survived the FDR-corrected permutation test, compared to 3243/4290 = 75.6% of between-participant correlations. Within the low-resolution atlas, 64/66 = 97.0% of within-participant correlations survived the permutation test, compared to 3029/4290 = 70.6% of between-participant correlations (Fig. S6A). Finally, we calculated individual-level identifiability, as the fraction of times that within-participant scan correlations are higher than between-participant scan correlations, using one of the contrasts as a reference (Fig. S6D). Individual identifiability was highly similar when using EPImix $T_1$-w scans and $T_1$-w scans as reference. Individual participants were most identifiable at the level of GM voxels, with high individual identifiability at the level of all brain voxels and regions of the high-resolution atlas as well; regions of the low-resolution atlas led to comparatively lower individual identifiability (Fig. S6D).

To dissect the effect of voxel-wise smoothing on across- and between-participant correspondence as well as identifiability, we repeated a subset of the above analyses after smoothing voxel-wise data using 2, 4, and 6 mm FWHM kernels, and compared results to unsmoothed data (0 mm FWHM below) (Fig. S7). We first inspected the correlation, across participants, of all brain voxels as a function of smoothing kernel size. The effect of smoothing was to reduce correlations; maximum correlations decreased as a function of smoothing, both within the whole-brain mask (max($\rho$) for: 0 / 2 / 4 / 6 mm FWHM = 0.79 / 0.78 / 0.71 / 0.60), and within the GM mask (max($\rho$) for: 0 / 2 / 4 / 6 mm FWHM = 0.70 / 0.69 / 0.61 / 0.55) (Fig. S7A). We next investigated between-participant correspondence using voxel-wise GM $T_1$-w intensities, which is the voxel-wise type of data for which identifiability was highest in unsmoothed data ($I_{diff} = 0.24$, compared to $I_{diff} = 0.12$ for all brain voxels). The effect of smoothing was to increase both within-participant and between-participant correlations, but with a greater increase in the latter; resulting in reduced differential identifiability as a function of increasing smoothing kernel size ($I_{diff}$ for: 0 / 2 / 4 / 6 mm FWHM = 0.24 / 0.24 / 0.21 / 0.16; Fig. S7B,C).

Finally, we constructed networks of $T_1$-w intensity covariance, using both EPImix and single-contrast scans. While single-contrast $T_1$-w structural covariance networks showed similar hallmarks of organisation to structural covariance networks commonly constructed from regional cortical thickness or grey matter volume data, such as strong long-range inter-hemispheric correlations between homotopic regions, structural covariance networks constructed from EPImix data instead showed high short-range correlations, clustered in frontal cortex; particularly so for regions of the high-resolution atlas (Fig. S8). The correspondence between the upper triangular parts of the structural covariance matrices was modest for the high-resolution atlas (Spearman's $\rho = 0.22$), with higher correspondence for the low-resolution atlas (Spearman's $\rho = 0.45$).

*Corresponding author
[1]These authors have contributed equally.

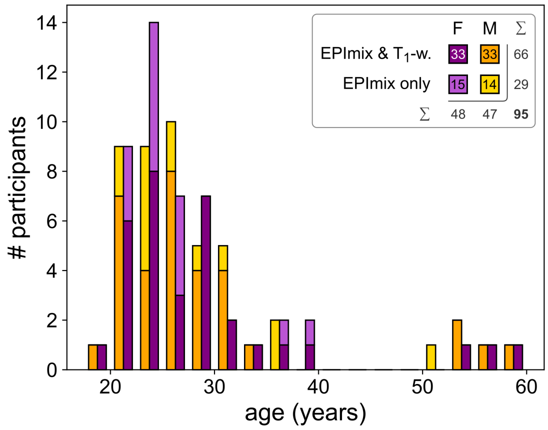


**Figure S1: Age distribution of participants by sex and scan sequence.** Scans from a total of 95 participants (48 female, 47 male) were included in this study. Of those, 66 (33 female, 33 male) were scanned using both EPImix and single-contrast $T_1$-weighted sequences, while an additional 29 (15 female, 14 male) were scanned using EPImix only. There were no significant differences in participant age by sex or scan sequence (Chi-squared test, $\chi^2 = 0.005$, P = 0.95).

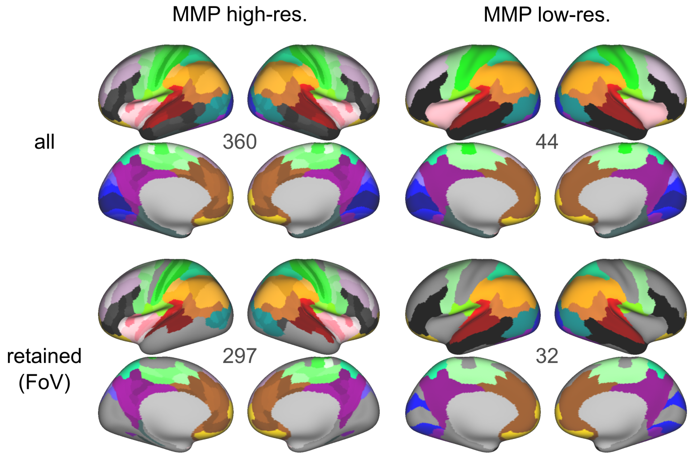


**Figure S2: Multi-modal parcellation (MMP) atlases used.** Top row: A multi-modal cortical atlas was used at two different spatial resolutions: a high-resolution version (Glasser et al., 2016) and a low-resolution version, created by downsampling the high-resolution atlas as described in Glasser et al., 2016 SI. Bottom row: Only regions with at least 80% EPImix scan coverage in at least 80% (76/95) participants were used for further analysis. Numbers within each panel correspond to the number of regions in each atlas version.

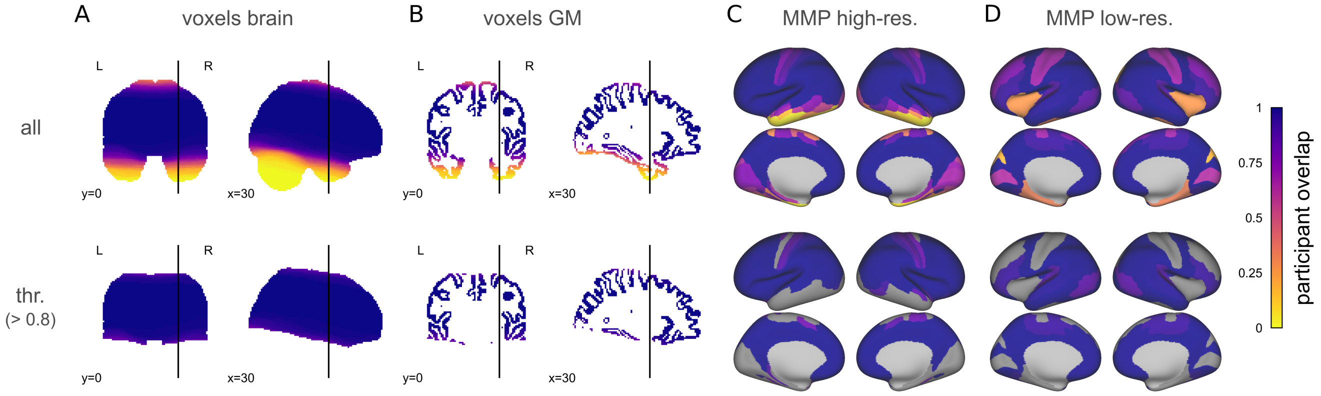


**Figure S3: Participant overlap at voxels and regions of interest in EPImix scans with reduced FoV.** Top row: Proportion of participants with data at each voxel of A) the (MNI) brain, and B) cortical grey matter (GM). The map in panel B (top) was used to calculate participant overlap within regions of interest, for both C) the high-resolution MMP atlas, and D) the low-resolution MMP atlas. In panels C and D, regions are color-coded by the proportion of participants with at least 80% (non-zero) voxels in each region. Subsequent analyses were limited to voxels and regions with at least 80% participant overlap (bottom row).

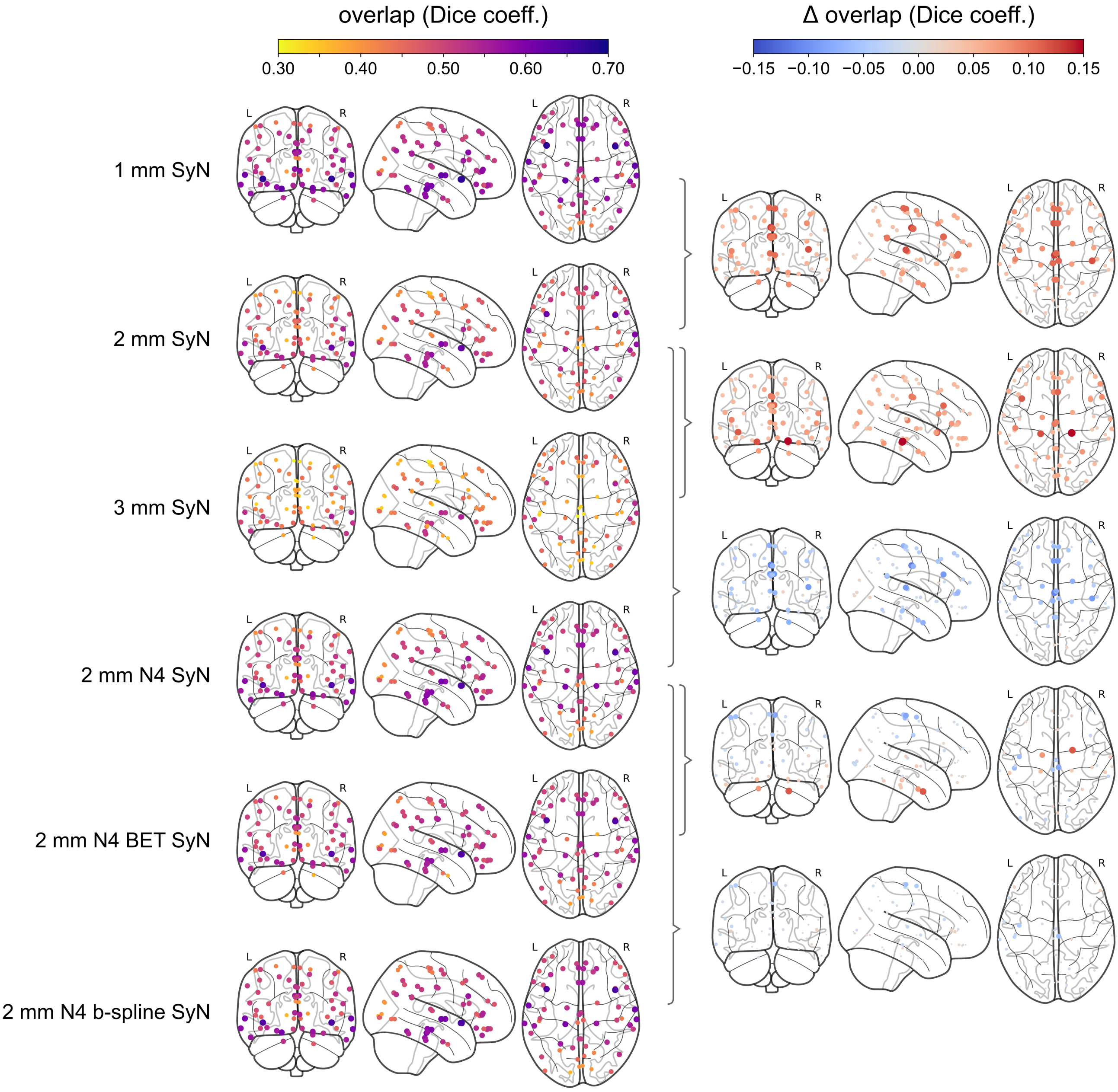


**Figure S4: Evaluation of regional quality of registration (and preceding steps) using the Mindboggle dataset.** Left: Regional Dice coefficient values quantifying the overlap between "manually" registered atlas labels and those released with the Mindboggle dataset (Klein and Tourville, 2012), for each of six evaluated processing pipelines (rows 1-3: spatial resolution; row 4: bias field correction; row 5: brain extraction; row 6: b-spline SyN registration). Right: Regional differences in Dice coefficient values. Pairs of maps being compared are joined by grey braces. Differences were calculated by subtracting values of the map below from the map above (i.e. $\Delta\text{Dice} = \text{Dice}_{above} - \text{Dice}_{below}$).

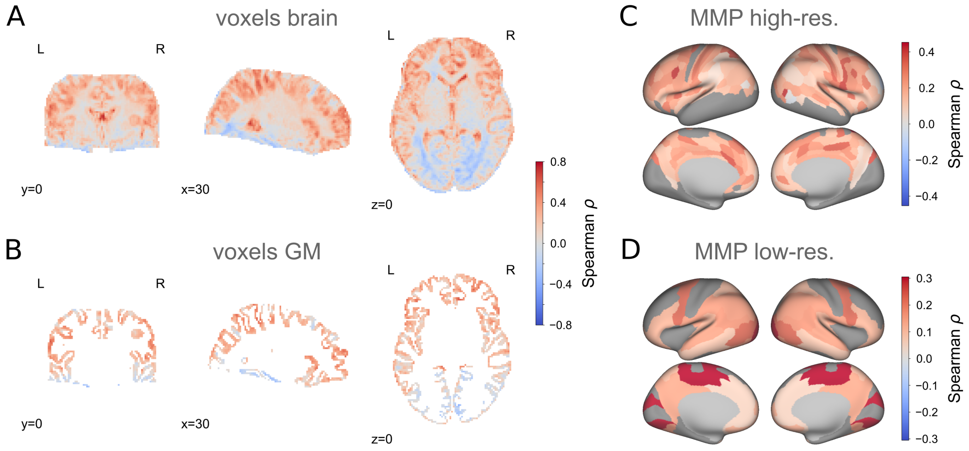


**Figure S5: Local correspondence of $T_1$-w intensities across participants.** Spearman's correlations between intensities of rapidly-processed $T_1$-w scans from the EPImix sequence and a single-contrast acquisition, using data of 66 participants. Correlations are depicted: at the voxel level for A) the whole brain, and B) cortical grey matter, as well as within regions of interest of C) the high-resolution and D) the low-resolution MMP atlas. (At the regional level, median regional values were extracted prior to calculation of correlations for each region.)

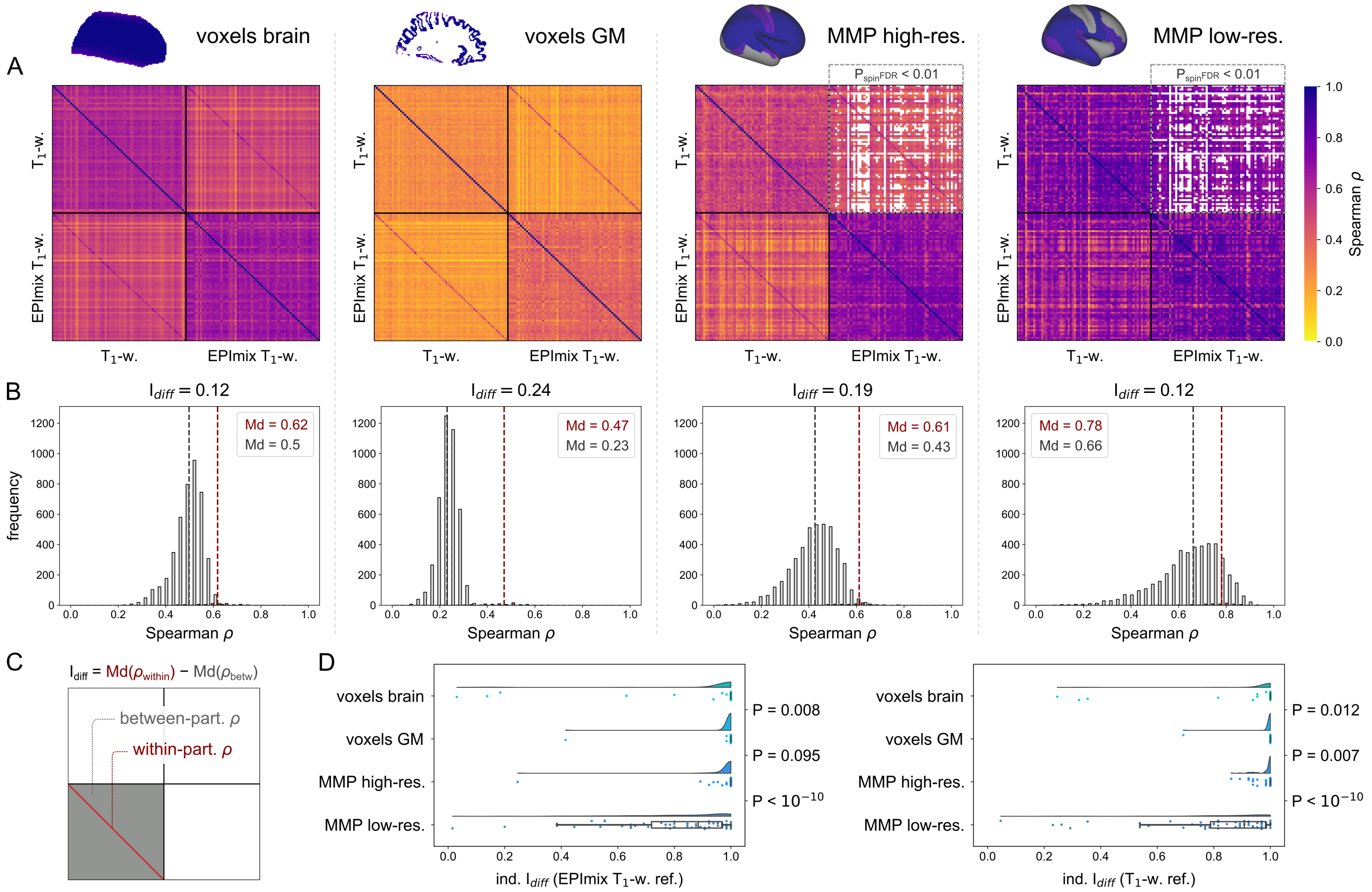


**Figure S6: Participant identifiability across EPImix and single-contrast scans, using $T_1$-w scan intensities.** Between-participant correlations and identifiability were investigated using four types of input data, at three spatial resolutions (columns in panels A-B, rows in panel D): all brain voxels, cortical grey matter (GM) voxels, regions of the high-resolution MMP atlas, and regions of the low-resolution MMP atlas. A) Spearman's correlations between EPImix and single-contrast $T_1$-w scan intensities, within and between participants. Cross-contrast correlations at the level of regions of interest were benchmarked using a null model controlling for contiguity and spatial autocorrelation (upper triangular blocks). B) Differential identifiability of contrasts, defined as the difference between the median within-participant correlation and the median between-participant correlation, as illustrated in C). D) Individual identifiability, defined as the fraction of times that the within-participant correlation is higher than between-participant correlations, either identifying a single-contrast $T_1$-w scan relative to EPImix $T_1$-w scans (EPImix $T_1$-w ref.), or vice-versa ($T_1$-w ref.). P-values correspond to the Wilcoxon rank-sum test between neighbouring distributions.

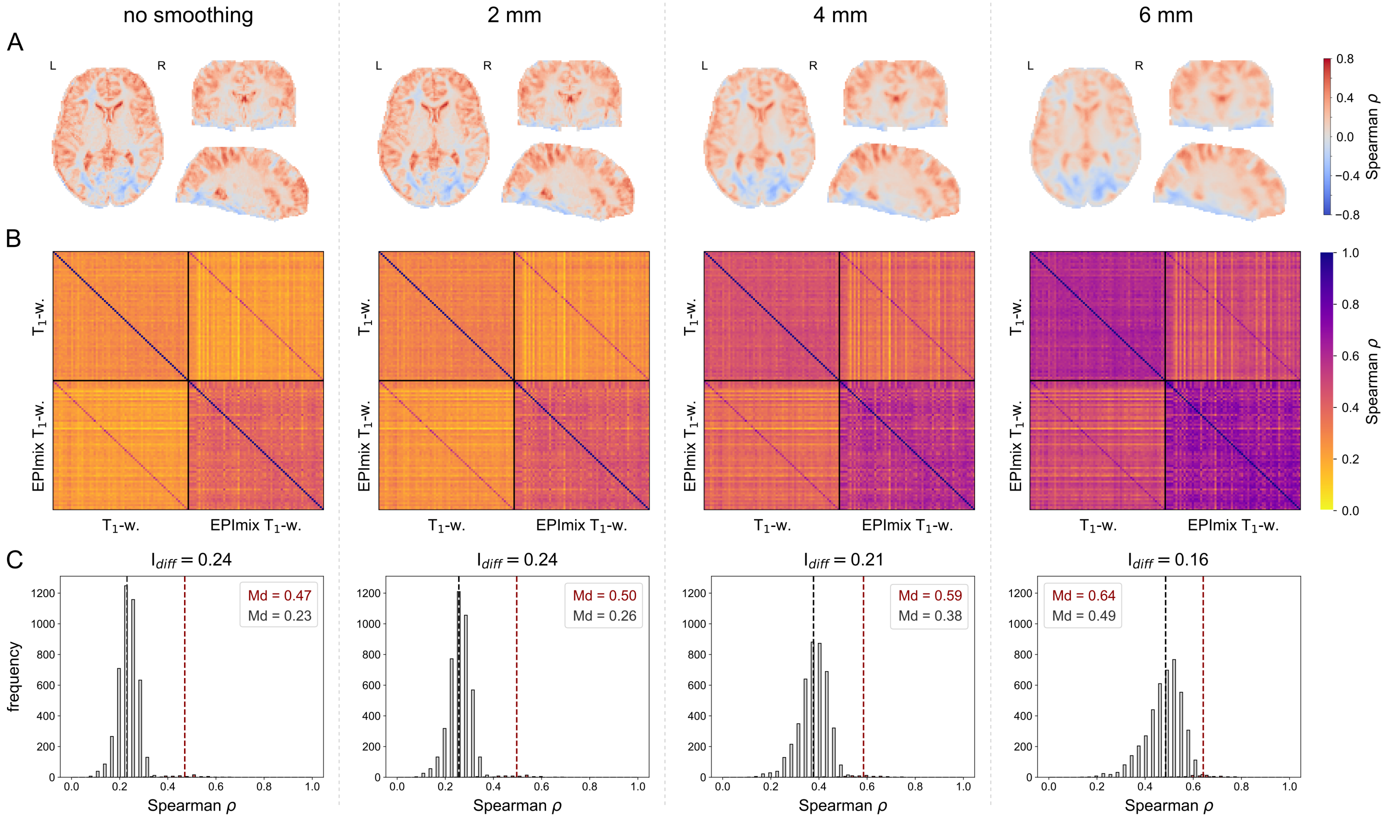


**Figure S7: Effects of data smoothing on between-participant correspondence and identifiability of voxel-wise GM $T_1$-w intensities.** A) Spearman's correlations between voxel-wise $T_1$-w intensities of rapidly-processed scans from the EPImix sequence and a single-contrast acquisition across 66 participants, as a function of smoothing. B) Spearman's correlations between EPImix and single-contrast $T_1$-w scan intensities, within and between participants. C) Differential identifiability of contrasts, defined as the difference between the median within-participant correlation and the median between-participant correlation (as illustrated in main text Fig. 5C).

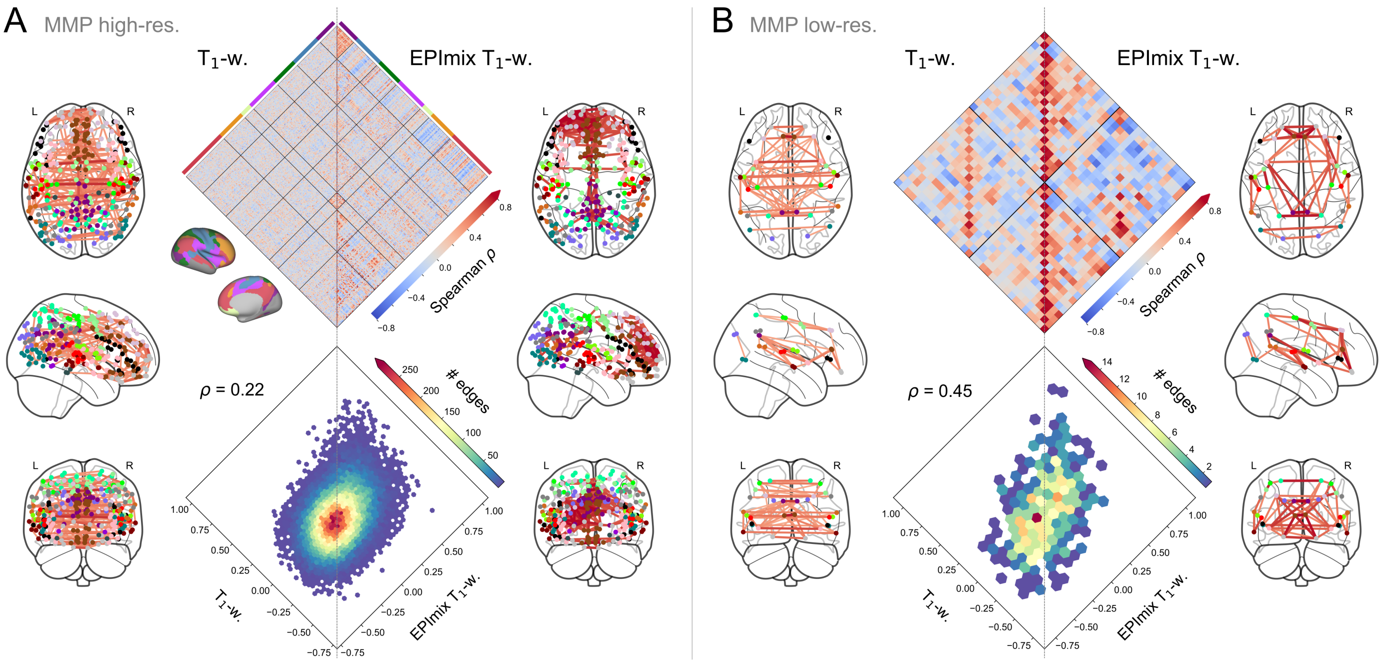


**Figure S8: Structural covariance networks constructed from EPImix and single-contrast $T_1$-w intensities.** A) Structural covariance networks constructed using the high-resolution MMP atlas (297 regions). The diamond plot (top) is ordered according to regional membership of the 7 canonical intrinsic connectivity networks derived by Yeo et al., 2011. Network diagrams depict the strongest 0.3% correlations. B) Structural covariance networks constructed using the low-resolution MMP atlas (32 regions). Network diagrams depict the strongest 10% correlations.
